# Phylofactorization - theory and challenges

**DOI:** 10.1101/196378

**Authors:** Alex D. Washburne

**Affiliations:** Montana State University

**Keywords:** compositional data, community ecology, isometric log-ratio, phylogeny, phylofactorization, graph-partitioning, greedy algorithm, regression, cross-validation

## Abstract

Data from biological communities are composed of species connected by the phylogeny. A greedy algorithm ‘phylofactorization’ - was developed to construct an isometric log-ratio transform whose balances correspond to edges along which traits arose, controlling for previously made inferences.

In this paper, the general theory of phylofactorization is presented as a graph-partitioning algorithm. A special case-regression phylofactorization-chooses coordinates based on sequential maximization of objective functions from regression on “contrast” variables such as an isometric log-ratio transform. The connections between regression phylofactorization and other methods is discussed, including matrix factorization, hierarchical regression, factor analysis and latent variable models. Open challenges in the statistical analysis of phylofactorization are presented, including criteria for choosing the number of factors and approximating null-distributions of commonly used test statistics and objective functions. As a graph-partitioning algorithm, cross-validation of phylo factorization across datasets requires graph-topological considerations, such as how to deal with novel nodes and edges and whether or not to control for partition order. Overcoming these challenges can accelerate our analysis of phylogenetically-structured data and allow annotations of edges in an online tree of life.

## 1 Introduction

It’s easy to take for granted the elegance of the particular spherical coordinates used to define locations on the surface of the Earth. First, the approximately constant radius of the Earth provides a natural reduction in the dimension of GPS data from 3 to 2 dimensions when changes in elevation are negligible. When changes in elevation are non-negligible, changes in radius correspond to changes in air temperature and atmospheric pressure independent of changes in the other two dimensions. Second, the choice of latitude to correspond to angular deviation from the equator in the direction of the axis about which the Earth spins may seem obvious, but on an abstract sphere one could choose any axis to define latitude. Latitude relating to the spin of the Earth about its axis-and closely corresponding to the revolution of the Earth about the sun-is a “ natural” choice of a variable that yields latitudinal associations with other important measurements such as climate (tropical, subtropical, temperate and arctic), positions of stars in the sky, and more. Longitude does not have an obvious reference, so it is set by convention to be zero at the Prime Meridian, which conveniently passes through the Royal Observatory in Greenwich, England (the French detested this convention, and used their own Paris Meridian until the early 20th century). Some coordinates are natural choices, whereas others are left to convention. If we didn’t know the Earth was a sphere and spinning about an axis, one would be pleased to discover two coordinates which so closely correspond to changes in important environmental meta data.

While distances often motivate coordinates for algebraic ease, the choice of coordinates is still separate from the choice of distances. One could define the same distance as the crow flies or “as the gopher burrows”-on the same sphere independently of the choice of coordinates. Thankfully, the distance between Sienna and Los Angeles remains the same, regardless what bizarre co-ordinates mathematicians are working with. The point being: choice of coordinates is separate from the choice of distance, and changing coordinates is motivated by dimensionality reduction, natural directions one might travel or along which meta-data change, and remaining coordinates can be filled in ad hoc, by convention, or for convenience.

Compositional data bound to the surface of the simplex yield many choices of distance, the most commonly used of which is the Aitchison distance [Aitchison, 1986]. Fixing the definition of distance between points, there are many choices of coordinates motivating more elegant, relevant, and easy-to-analyze quantities for data on the simplex. The isometric log-ratio (ilr) transform [Egozcue et al., 2003, Egozcue and Pawlowsky-Glahn, 2005] is a change of basis to variables whose Euclidean distances are equal to Aitchison distances of the original, compositional variables, and whose values reflect a contrast or the differences in relative abundance of two groups. The ilr transform can be defined through a sequential binary partition defining which groups of parts to contrast for each coordinate but, in choosing which isometric log-ratio transform to use for analysis and cross validation across datasets, one encounters similar considerations as those encountered when choosing spherical coordinates for the Earth [Pawlowsky Glahn and Buccianti, 2011].

This paper is about a method-phylofactorization [Washburne et al., 2017]-whose aim is to construct an isometric log-ratio transform for biological data which yields coordinates that are consistent with the compositional nature of the data, meaningful (biologically interpretable), and convenient (coordinates along which there are predictable changes). Much like spherical coordinates for GPS data require geography and astronomy to motivate, phylofactorization requires some biological background to motivate the coordinates and precisely why we are choosing them.

First, one needs justification for the analysis of communities as compositions or, generally, as objects prone to geometric changes and thus justifiably analyzed by isometric log-ratios. Then, one needs to know about the tree of life (the phylogeny) as a graph with no cycles, a sequential binary partition connecting species in a community. Phylofactorziation often uses the ilr transform as a contrast of two groups separated by edges in the phylogeny. Variables corresponding to edges in the phylogeny yield biologically meaningful coordinates corresponding to putative functional ecological traits, axes about which biologists might expect there to be predictable changes with environmental meta-data, which may allow further development of community ecological theory [McGill et al., 2006].

This paper discusses the general theory of phylofactorization as a graph-partitioning algorithm iteratively cutting the phylogeny at edges which maximize an objective function. Edges in the phylogeny separate the community into two disjoint groups of species, and thus, when analyzing community composition, one can use ilr coordinates as quantities of interest corresponding to edges. Phylofactorization done to maximize objective functions from regression on ilr coordinates is called “regression-phylofactorization”; the relationship between regression phylofactorization and matrix factorization, hierarchical regression, factor analysis, and latent variable models is discussed. Future research directions are discussed, including criteria for choosing the number of factors, the null distribution of test-statistics from phylofactorization to allow hypothesis-testing, and graph-topological considerations when cross-validating phylofactorization across datasets with non-identical, or even disjoint, sets of species. Future research along these directions carries major implications for methods to diagnose disease [Kostic et al., 2014], modulate microbial communities [Rajpal and Brown, 2013], and make inferences about the habitat associations of unclassified species in the tree of life [Letunic and Bork, 2007].

## 2 Communities as Compositions

Communities are assemblages of organisms in a study area or a sampling design. The definition of a community is often arbitrary and based on the organisms we happen to observe or we think are important. Due to the challenge of sampling communities, communities are rarely the entire set of species in a region of space-rather, communities will be defined as the assemblage of trees in a forest, grasses in a grassland, mammals in Yellowstone National Park, or birds in a patch of rainforest. For example, one could analyze the community of non-human animals around Sienna. If a researcher counted the non-human animals around Sienna, they would observe counts, integer valued random variables of the numbers of birds, cows, sheep, cats, dogs, and other non-human animals in the community.

Community ecology studies how communities assemble, function, and respond in light of biotic and abiotic conditions. A common question is: how do communities differ along environmental gradients or between sample sites? One of the most famous and pionerring examples of this line of questioning was Alexander von Humbolt’s observation that how plant communities change with increasing latitude is very similar to how plant communities change with increasing altitude, implicating climate (rainfall, length of growing season, etc.) as an important abiotic factor in community structure and function. For an example closer to CoDa 2017, how would the animal community in Sienna differ from the animal community in the Mediterranean Sea, and how much of that is due to latitude (likely very little) versus the fact that one habitat is terrestrial and one is marine? We obtain count data of organisms, and wish to make inferences on how these counts change and which environmental meta-data are most important.

Compositional data analysis has a long marriage with geosciences, but it is somewhat unfamiliar to community ecology, and so it’s worthwhile to take a few paragraphs to motivate compositional data analysis in community ecology. The first step in this paper is to motivate communities as compositions or, more generally, as comprised of quantities which exhibit geometric changes most appropriately analyzed in terms of the log ratios in compositional data analysis. Once we believe communities are best analyzed in terms of log ratios, we can move on confidently to discuss the ilr transform of a community as a means of analyzing problems like the von Humbolt’s, and then we can discuss how the ilr transform allows us to choose very meaningful coordinates in light of the evolutionary tree.

First, consider the nature of the data one obtains when sampling communities. Suppose we have a small sample size where we observe 10 doves, 30 starlings, and 10 cats in one day, and, on a rainy day, we observe 2 doves, 6 starlings, and 2 cats. Did the population of doves decrease by 8, or did our sampling effort/ability change? Many animals are not as active on rainy days, so one would clearly attribute this to a change in effort or detectability. To control for changing effort or detectability, a more robust analysis would limit itself to inferences on the relative abundances of doves, starlings, and cats, possibly incorporating sampling effort/ability as weights for averages, regression, etc.

Are communities only compositions when we obtain a small sample size? What if we sampled every non-human animal in Sienna, and the total community size changes (e.g it grows as people acquire fewer cats and fewer birds are killed, or shrinks in a drought)? In small islands, fragments of habitat or enclosures, or extensively sampled habitats with easily detectable organisms, one knows absolute abundances how should we analyze changes in absolute abundances?

Biologists have long struggled to define first principles, but I will propose two such principles central to the definition of “life”-reproduction and death. Every organism can die, and just about every organism can reproduce. If there was no reproduction and every organism had a constant probability of dying per unit time, the expected population dynamics would be an exponential decrease in population size, or linear decreases in log-abundances over time. If there is reproduction, and the propensity for birth exceeds that of death, the expected population dynamics would be an exponential increase in population size, or linear increases in log abundances. More complicated state-dependent propensities can yield complicated dynamical systems, yet, near an equilibrium where birth rates equal to death rates, all such dynamical systems exhibit exponential approach to or explosion away from equilibria, most conveniently analyzed as changes in log abundances. If changes in environmental conditions lead to changes in fitness, i.e. per-capita propensities for reproduction and death, we would see short-term, geometric changes in population size. Motivated by these first principles of percapita birth and death yielding a tendency for exponential growth and decay, one is not without justification to default to analyzing absolute abundances on a logscale, with changes modeled as log-ratios. There is empirical support for analyzing population dynamics on a log-scale: Kalyuzhny et al. [2014] looked at bird populations across North America from 1966 to 2014 and found that the counts of hundreds of sub populations of birds exhibit temporal fluctuations more consistent with a geometric Brownian motion than by an arithmetic Brownian motion, more naturally and coherently analyzed in terms of changing log ratios of abundances. Populations move more like a stock price than a particle under a microscope.

So, if we count only a few birds, we are best confined to using log ratios, with appropriate treatment of zeros, to infer changes in relative abundances. If we count all the birds, we are at least somewhat justified in using log-ratios to infer changes in absolute abundance. Changes in equilibrium population sizes and community compositions may be neither arithmetic nor geometric, as changes in rainfall can lead to complicated, neither linear nor exponential, increases in the number of trees in a region, so there is no law that log ratios are always appropriatethey should be justified in each case. However, most datasets do not sample entire communities and so, often, changes in community composition are justifiably analyzed as changes in log-ratios.

Most phylofactorizations of community ecological data have focused on a recent and highly relevant class of data-“microbiome” data, collected by sequencing microbes in a region such as soils, tongues, or the human gut. These data are obtained by amplicon sequencing-sequencing “barcode” genes and counting the number of different types of “barcodes”-obtained where absolute counts depend on the amount of reagents we put into a machine (the sequencing depth) and not absolute abundances of microbes in the community. These sequence-count data are as compositional as the bird counts in Sienna-the absolute number of counts relates to effort, and only inferences on relative abundances can be made. While these data are compositional, best analyzed with log-ratios and Aitchison distances, the choice of coordinates remains: changes are relative, but which groups are changing, and relative to which other groups are they changing? Microbiome datasets often come with the evolutionary tree articulating the common ancestry and origins of species in the dataset, a natural scaffolding for choosing groups and, therefore, choosing coordinates.

## 3 The Evolutionary Tree

All cellular life arose from a common ancestor, and so all community ecological datasets contain parts connected by the evolutionary tree also known as the phylogeny. The phylogeny simplifies biological data into a nested hierarchy of lineages-doves are birds, birds are vertebrates, vertebrates are animals. Going the other direction, animals can be split into vertebrates and invertebrates, vertebrates can be split into amniotes and anamniotes, and so on to the doves, starlings and cats found in Sienna.

The evolutionary tree contains a natural means of dimensionality reduction in biological data. Consider describing the 66,000 vertebrate species based on whether they live on land or in water. One could capture all of the land/water associations with 66,000 variables, one for each species. Alternatively, one could note that all vertebrate lineages before tetrapods (e.g. fish, sharks, rays) live in the water, whereas tetrapods tend to live on land. One variable indicating whether a species is a tetrapod or not, a variable corresponding to one edge in the phylogeny, captures most of the 66,000 species’ land/water associations by accurately guessing the habitat of over 30,000 fish, sharks and rays and correctly guessing the habitats of most reptiles, birds and mammals. A handful other, similarly constructed variables will finish the job: whales and dolphins live in the water relative to other tetrapods (a second variable splitting whales & dolphins from all other tetrapods), seals/sea lions/walruses live in the water (a third variable splitting seals/sea-lions/walruses from all tetrapods that are not whales/dolphins), and so on (partitioning amphibians that live on land or in the water will likely require some more variables). A handful of well-chosen coordinates corresponding to edges on the tree of life can explain most of the variance in land/water associations in a 66,000 dimensional dataset.

The utility of the phylogeny for dimensionality reduction stems from its correspondence to traits, the products of evolution by natural selection, features which determine where an organism lives, what it eats and how it responds to changing environmental conditions. The edge along which tetrapods arose is the edge along which limbs and lungs arose, allowing organisms to walk on land and breath air. The edge separating whales & dolphins from their most recent common ancestor on land is the edge along which flippers and blowholes arose, allowing them to swim and conveniently gasp for air at the surface of the water. While we’re more familiar with the traits of animal lineages and their ecological functions, we know very little about the traits on the microbial tree of life. For microbes, we have a phylogeny, often constructed from the “barcodes” mentioned above, but we know almost nothing about which traits arose on which lineages and what functions those unknown traits may have. The study of microbial communities can be improved by considering traits, either explicitly as measured or implicitly as latent variables on the tree of life, as determinants of disease, habitat association, or response to perturbation [Martiny et al., 2015].

We’ll be dealing with the phylogeny in this paper as a mathematical structure used to make inferences about locations of putative traits driving changes in community composition. As a mathematical structure, the phylogeny is a connected graph containing no cycles. For a given gene, the true structure is a sequential binary partition-a strictly bifurcating graph where all internal nodes have three neighbors and all tips have one. However, phylogenies are never known-they are estimated-and often we don’t know how to resolve the relationships between ancient nodes as no one was there to see what happened. We use contemporary data to guess what might have happened. Unresolved nodes are “polytomies”, and a tree with polytomies is not a sequential binary partition; an unresolved polytomy will have one parent and more than two descendants. All edges, however, connect only two nodes. The edges in the phylogeny have lengths, referred to as “branch lengths”, which approximately correspond to the time between speciation events. Edges are the locations along which traits arise.

## 4 Phylofactorization

If we observe changes in the composition of a community of species connected by an evolutionary tree, which coordinates best capture the changes and relate to putative traits?

Consider a simulated dataset of community compositions, ***x***_*j*_*∈* Δ^*D*^ for *D* = 10, across 10 samples *j* = 1*, …,* 10, illustrated in Figure 1. The dataset consists of a set of vertebrates one might find in Australia sampled in 5 different sites in each of two different environments. In these data, we consider traits driving habitat associations. Traits arise along edges and, in the example in Figure 1, two traits drive differential abundances. One trait, “A”, arose along the common ancestor shared by hawks and owls and leads to an increased abundance in environment B. Another trait, ‘B’, is shared by mammals and leads to an increased abundance in environment B. A default analysis of community compositions would motivate using a clr or an ilr transform.

**Figure 1:**
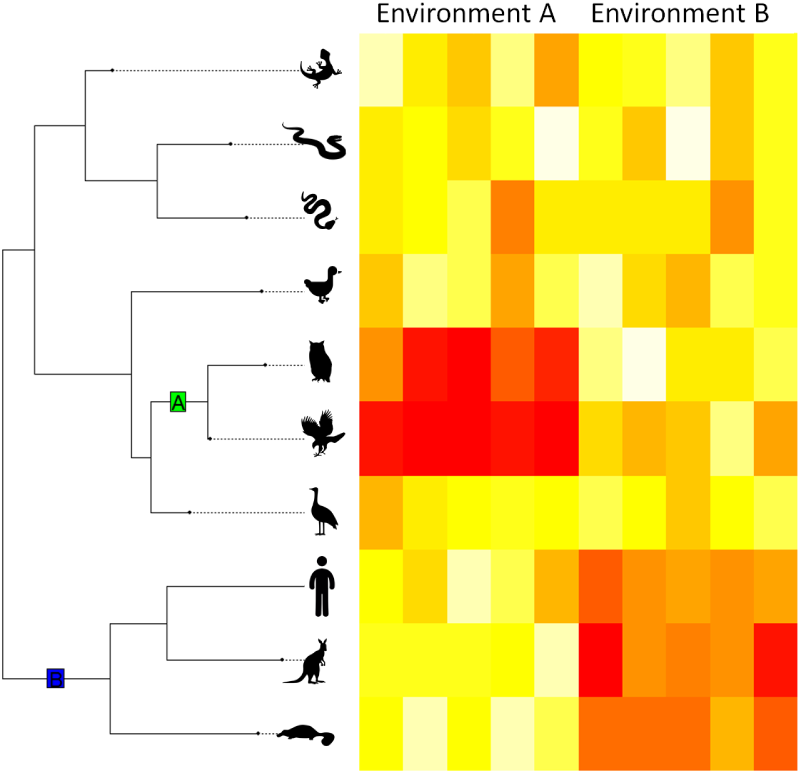
Phylogenetically structured compositional data. A simulated dataset of relative abundances of 10 species across 5 samples in each of two environments. Blocks of data correspond to the phylogenetic structure one would observe if monophyletic clades have common patterns of abundance across environments. In this example, raptors (owls & hawks) are hyper abundant in Environment A, whereas mammals (humans, kangaroos and platypus) are hyper abundant in Environment B. Common habitat associations of organisms with common ancestry may be caused by functional ecological traits.

Analysis of an arbitrary ilr transform may indicate differences between the two environments, but arbitrary ilr coordinates do not map to traits. We can do better. The phylogeny is a sequential binary partition, but analysis of an ilr transform constructed from the rooted phylogeny will obtain variables corresponding to changes in sister clades-such as changes in hawks and owls relative to changes in cranes. The inelegance of the ilr transform of a rooted phylogeny comes from the variables corresponding to nodes, not edges. The changes we aim to detect are not changes of sister clades relative to one another, but changes in one clade with a trait relative to the rest of the organisms without a trait, possibly controlling for other traits we have already identified as important. Such changes correspond to edges, giving us coordinates that can be interpreted as differential abundances of species with and without various traits. Constructing ilr coordinates corresponding to edges yields coordinates corresponding to latent variables-traits. However, the edges don’t define a sequential binary partition-we need to choose one.

### 4.1 General algorithm for phylofactorization: a graph-partitioning algorithm without a balance constraint

Phylofactorization [Washburne et al., 2017] is a greedy algorithm for constructing a sequential binary partition corresponding to edges in the phylogeny. Each edge, *e*, in the phylogeny separates the community into two disjoint groups of species, *R*_*e*_ and *S*_*e*_, containing *r*_*e*_ and *s*_*e*_ species, respec tively. Phylofactorization requires an objective function, *ω*(***X****, R, S*), of the dataset, ***X*** = (*x*_*i,j*_) and the disjoint index sets of the two groups, *R* and *S*. The most general form of phylofactorization defines a graph-partitioning algorithm [Buluç et al., 2016].

The general algorithm for phylofactorization follows:

1. Compute objective function corresponding to each edge, *e*, with partition *p_e_* separating two groups, 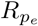 and 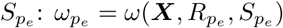

2. Identify 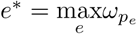

3. Cut the tree at *e*^*\**^, creating two disconnected graphs

4. Repeat 1-4 until stopping criterion is reached.

The general algorithm is illustrated in Figure 2.

**Figure 2:**
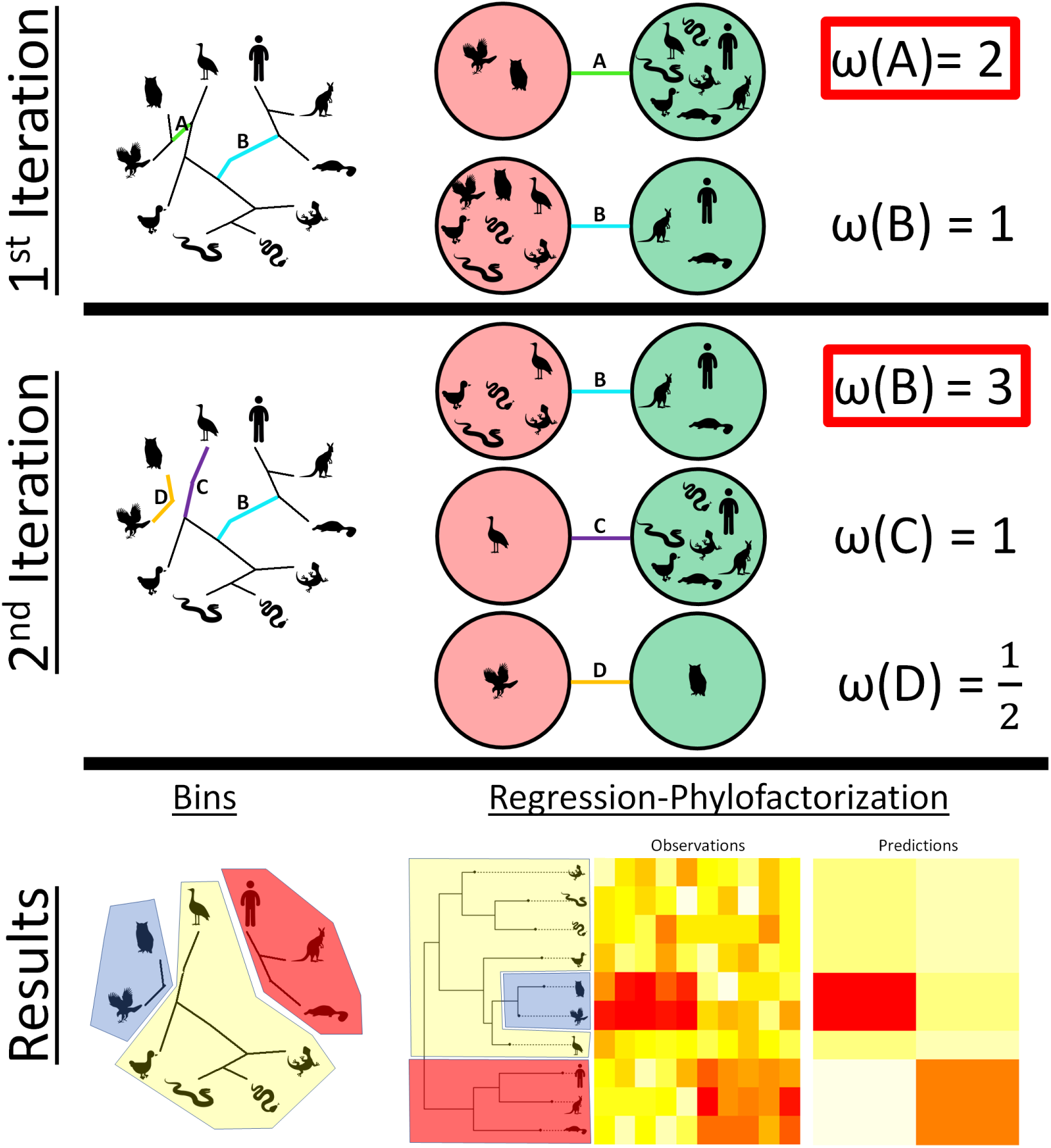
Phylofactorization. Phylofactorization is a graph-partitioning algorithm which iteratively cuts edges in the phylogenetic tree based on objective functions, *ω*. In the first iteration (**top row**), all edges are considered, but we depict the two edges with traits affecting abundance patterns in figure 1. Each edge separates the group of *D* species into two groups. Edge A separates the raptors (hawk and owl) from all other species, and edge B separates the mammals (platypus, kangaroo, and human) from all other species. An objective function, *ω*, often a measure of contrast of the two groups is calculated for each edge, and the edge which maximizes the objective function is selected. The phylogeny is cut/partitioned at that edge-edge A, in the illustrated example-and the algorithm continues. In the second iteration (**middle row**), all edges are considered but each edge separates only those species in its sub-graph. Since edge A was cut in the first iteration, in the second iteration edge B contrasts mammals from all non-raptor species, not mammals from all other species as it did in the first iteration. The edge which maximizes the objective function, in this case edge B, is chosen and the tree is cut. (**bottom row**) After *k* iterations of phylofactorization, one has identified *k* edges or chain of edges (see chain labeled edge D) of interest which partition the phylogeny into *k* + 1 bins of species. In the special case of regression-phylofactorization, edges yield contrasts between groups via projection of log relative abundances or absolute abundances onto a balancing element, *𝒱p*, corresponding to the partition, *p*, defined by the edge, and the objective functions are statistics from regression on the balance. Regression phylofactorization outputs predictions of *k* balances yielding low rank predictions of abundance patterrns, recreating the major blocks of variation illustrated here.

For example, one could define the objective function as a function of an ilr balance for each edge. Recall the ilr transform of a given sample corresponding to a generic partition, *p*, of two disjoint index sets, *R* and *S*, containing *r* and *s* species, respectively, can be written as

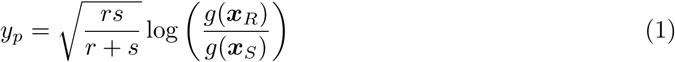

where *g*(.) is the geometric mean, 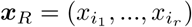 for all elements *i*_·_ *∈ R* and 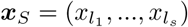 for all elements *l*_·_ *∈ S*. The balance, *y*_*p*_, can also be obtained by projecting log relative abundances, log(***x***), onto the balancing element, *𝒱*_*p*_, whose *i*th element is

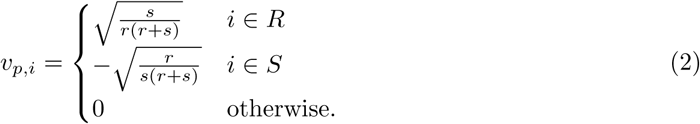

For each sample, *j*, one obtains *y*_*p,j*_ and can define the objective function as the variance of an ilr transform:

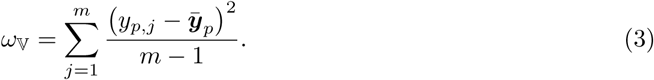

Maximizing *ω*_𝕍_ is similar to a principal components analysis-the edge, *e*^*\**^, with corresponding partition, *p*^*\**^, which maximizes *ω*_𝕍_ defines an ilr balancing element, *v*_*p\**_, onto which projection of log relative abundances maximizes variance. Phylofactorization by *ω*_𝕍_ may yield edges separating the most different species, corresponding to the most important traits in the dataset which may or may not be predictable with existing meta data. One could devise other objective functions with ilr balances, such as 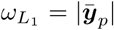. The balancing elements in the ilr transform are convenient tools for phylofactorization, allowing edges to be boiled down into variables interpretable as contrasts of groups split by the edge. The variance of the contrasts from projection onto balancing elements does not change with *r* or *s*, allowing a more fair competition of the edges in the phylogeny to be the site of a partition.

As a graph-partitioning algorithm, phylofactorization has dual interpretations in terms of the edges cut and the sub graphs that remain un cut. After *k* iterations of phylofactorization, the species will be split into *k* + 1 groups or “bins”, referred to as “binned phylogenetic units”. Where phylofactorization correctly identifies edges corresponding to functional ecological traits, the bins correspond to species with similar functional ecological traits. For instance, splitting vertebrates along the three edges: “tetrapod”, “Cetacean” (whale/dolphin), and “Pinniped” (seals, sea lions & walruses), will result in four groups: (1) non tetrapod vertebrates, (2) Cetaceans, (3) Pinnipeds,(4) tetrapods which are neither Cetaceans nor Pinnipeds. Group (1) has gills and fins and lives in the water, group (2) has flippers and blowholes and lives in the water, group (3) has flippers and eats fish and lives in the water, and group (4) has legs, lungs and lives on land. Phylofactorization can be seen as having two complimentary goals: accurately identifying edges corresponding to functional ecological traits, and accurately binning species into groups with similar traits. Novel objective functions or phylofactorization algorithms can be evaluated by their ability to perform each of these two goals.

In the general theory of phylofactorization, there are many unanswered questions for future research. The full connections to graph-partitioning including the biases of common objective functions for choosign basal or distal edges, the utility of different objective functions for inferring different traits and ecological processes has been largely unexplored, and the performance of Hastie Tibshirani [Friedman et al., 2001] stochastic algorithms may yield more accurate inferences but has yet to be investigated.

### 4.2 Regression phylofactorization

Often, researchers are interested how communities change across sample sites or environmental gradients such as latitude, pH in soils, or rainfall. For such problems, each community composition, ***x***_*j*_, comes with *m* associated environmental meta-data measurements, ***z***_*j*_= (*z*_1*,j*_, …, *z*_*m,j*_). Phylo factorization can be utilized by defining objective functions from regression on an ilr balance, such as the explained variance, *F*-statistics, or *t*-statistics from a focal coefficient in multiple regression. For compositional data, regression-phylofactorization performs regression on the ilr balances, 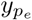 corresponding to the partition, *p*_*e*_, for each edge, *e*. The regression takes the form

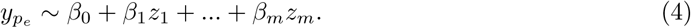

and one selects the edge, *e*^*\**^, with partition *p*_*e\**_, which maximizes the objective function of re gression. Regression phylofactorization is a form of hierarchical regression in which each iteration considers candidate variables,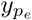, to maximize an objective function from regression and successive iterations consider variables with non-overlapping partitions, thereby controlling for previous inferences. However, instead of a strict order for hierarchical regression as used in ordinary least squares, phylofactorization hosts a competition at each iteration, *k*, among a set of candidate variables.

The objective function originally considered, and commonly used in exploratory regression phylofactorization, is the explained sum of squares from regression,

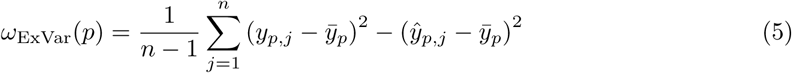

where 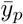 is the sample mean of ilr balance corresponding to partition, *p*, and 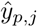 is the fitted value of regression on the ilr balance, *yp*. The use of explained sum of squares as an objective function was motivated by the fixed total variance in a compositional dataset, irrespective of the choice of ilr transform, which implies that each iteration of phylofactorization has candidate edges, *e*, competing for a fixed amount of remaining variance to explain. Explicitly,

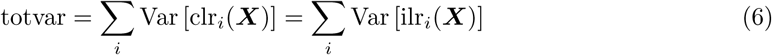

for any choice of ilr transform. After *k* iterations, the previously chosen ilr balances with partitions 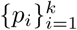 will explain a sub total variance,

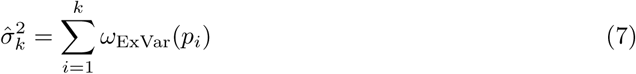

leaving candidate ilr balances competing for totvar - 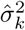variance in the dataset. Maximizing the explained variance from regression at each iteration of phylofactorization produces a set of ilr balancing elements with fitted values that provide a low-rank approximation of a dataset, clr(***X***), discussed further in the next section: Phylofactorization and factor analysis.

Regression phylofactorization can also be used in sample site classification, a crucial problem for disease detection, by reversing the regression:

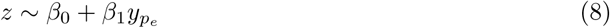

and maximizing the *F* statistic to identify phylogenetic bioindicators of disease. Ideally, one would like to use multiple regression to produce multiple bioindicators of disease, but the power of such approach is limited by the sequential nature of phylofactorization. The computational time needed to perform multiple regression of the form

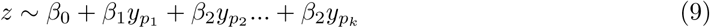

grows exponentially with *k*, and many problems of partitioning a graph into *k* bins are NP hard. However, solutions or algorithms which perform reasonably well at finding the set of partitions which maximize the *F*-statistic in the regression of Equation (9) may improve the classification of disease, such as inflammatory bowel disease [Kostic et al., 2014], and identification of microbial drivers of environmental conditions.

### 4.3 Phylofactorization and factor analysis

Recall that the ilr transform in Equation (1) can be re written as a projection of log relative abundances, log(***x***), onto a basis vector called the balancing element, *𝒱*, defined in Equation (2). In *K* iterations of regression phylofactorization, one has a set of balancing elements, 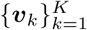, giving a set of *K* balances, 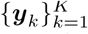, and their fitted values from regression,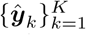. Let *V* be a *D × K* matrix whose columns are balancing elements, *𝒱*_*k*_, *Y* the *K × n* matrix whose rows are ilr balances, *y*_*k*_ corresponding to the respective balancing elements in the columns of *V*, and whose columns are samples. Let *Ŷ* the same as *Y* but containing fitted balances, 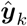. Regression phylofactorization can serve as a rank *K* approximation of the clr-transformed data matrix

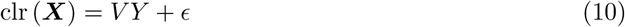

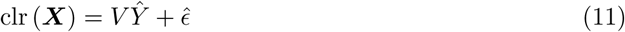

where *ϵ ∈* ℝ^*D×n*^ is the error of the approximation by ilr balances and 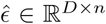the error of the approximation by fitted ilr balances. The matrix illustrated in the end of Figure 2 is an example of Equation (11), a rank 2 approximation of clr(*X*) using the fitted ilr balances from regression. In this sense, phylofactorization is a change of basis and a matrix factorization algorithm.

In the original paper, phylofactorization was also hypothesized to be a form of constrained factor analysis of the clr transformed data, as the use of ilr balancing elements allows the construction of a set of orthonormal axes that allow low rank predictions of the data. Factor analysis obtains low rank approximations of a data matrix, *Z ∈* ℝ^*D×n*^ through the product of an orthogonal matrix of “loadings”, *L ∈* ℝ^*D×K*^ and a matrix of “factors” *G ∈* ℝ^*K×n*^, plus an error matrix *ϵ*_*z*_*∈* ℝ^*D×n*^,

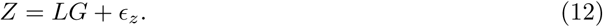

such that (1) *G* and *Ψ* are independent, (2) 𝔼[*G*] = 0, and (3) Cov[*G*] = *I*. Phylofactorization was hypothesized to be a form of factor analysis due to its apparent similarity in constructing an orthogonal matrix, *V*, a matrix *Y* corresponding to a latent variable (differential abundance of traits) and constrained to axes which are balancing elements of edges in the phylogeny.

However, the claim that phylofactorization is a form of factor analysis is not true for the default regression-phylofactorization maximizing *ω*_ExVar_. While phylofactorization clearly obtains a low rank approximation nominally similar to factor analysis, it is clear that 𝔼[*Y*] is not necessarily zero there could be imbalances in the data such that one group, *R* is, on average, more abundant than its complement, *S*, causing *g*(***x***_*R*_) *> g*(***x***_*S*_) and 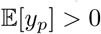for a given row of *Y*. Furthermore, while it has been conjectured that the factors are sequentially independent, under the assumption that an ilr balance at one partition contains no information about the ilr balance at subsequent partitions (the converse is not necessarily true: ilr coordinates along the root path of a sequential binary partition may be correlated under increases in single clades, as highlighted in the original phylofactorization paper), the conjectured independence of sequential factors (1) is a conjecture and (2) does not imply that Cov[*Y*] = *I* for a given phylofactorization. Whether or not sequential factors are independent depends on the underlying statistical model of how abundances change and whether or not a factor was correctly identified. There may be other algorithms for regression-phylofactorization which are a form of factor analysis, such as phylofactorization of standardized datasets (thereby assisting 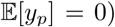 with objective functions which minimize off-diagonal elements of Cov[*Y*] or yield Cov[*Y*] = *I* asymptotically..

Despite phylofactorization not being factor analysis *sensu stricto*, there are conceptual similarities between phylofactorization, factor analysis, principal components analysis, and other methods. Such connections may carry implications or suggested solutions for many of the challenges highlighted in the next section. The orthonormal basis 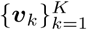chosen by phylofactorization is a low-rank set of orthogonal axes along which meaningful change occurs in a compositional dataset.Where the objective function is to maximize the variance of the ilr balance, phylofactorization becomes like PCA in finding orthonormal axes sequentially maximizing the variance of data projected onto the axis. Since the ilr transforms are all rotations of one-another, there may be connections with rotation methods in factor analysis to allow for convenient interpretations of the method. For instance, the bins described in the results of Figure 2 can be constructed either by factoring as illustrated (edge A and then edge B), or by a re ordering of the factorization of those same edges (edge B and then edge A)-the ilr bases corresponding to those two examples are rotations of one another. Finally, the “factors” in factor analysis are latent variables, and regression phylo factorization is inferring latent variables-traits and their differential abundance-that underlie covariances among species. In this sense phylofactorization is also a latent variable model, and could be implemented with more complicated functional responses of traits than those captured by regression on ilr balances. Phylofactorization also constructs a decision tree for the classification of species [Rokach and Maimon, 2014]-given a new species, we ask a set of questions to make estimates about its habitat associations (Is it a tetrapod? If yes, is it a whale/dolphin? If no, is it a seal/sea lion/walrus?).

The place of phylofactorization in the mathematical, statistical and computational literature, and its connections with other methods is still being resolved and may likely vary depending on the choice of objective function and particularities of the analysis. Future work integrating phylo factorization into more general mathematical and statistical literature may provide theoretical connections which suggest more efficient algorithms, more powerful tests, more apposite objective functions, and more interpretable results.

## 5 Challenges of Phylofactorization

The novelty and biological utility of phylofactorization leaves open many avenues of future research. As articulated above, more needs to be done to understand the generality, utility, and limitations of phylofactorization. In addition to articulating the place of phylofactorization in graph-partitioning algorithms, understanding the pros, cons and interpretations of different objective functions for regression phylofactorization, and understanding the relationship between phylofactorization and other statistical methods, there are several more focused challenges to the maturation of phylo factorization which, if addressed, can dramatically improve its utility in the biological sciences. Two of those challenges are illustrated here: criteria for choosing the number of factors and cross validation.

Much like factor analysis, there is the need for criteria to choose the number of factors, a problem intimately tied with the null distributions of test statistics under phylofactorization. At which iteration, *K*, should we stop factorization to control our family wise error rate of inferences on edges or control error rates in some other sense? Calculation of the Marchenko Pastur distribution [Marchenko and Pastur, 1967] for the asymptotic distributions of singular values aided the development of PCA into an inferential tool-what is the null distribution of the variance of the dominant edge’s ilr balance from phylofactorization with *ω*_𝕍_? What is the null distribution of the F-statistics from regression-phylofactorization? Answering these questions can move phylofactorization from an exploratory to an inferential tool.

Phylofactorization makes predictions assignable to a universal tree of life, which allows comparison of datasets with unrelated species for cross validation of phylofactorization. If we discover the ratio of birds to mammals shifts dramatically across two Australian habitats-tree tops and the desert-we may be interested in cross-validating such findings for birds and mammals on other continents, even if the sets of species across continents are completely disjoint. Cross-validation introduces graph topological and statistical challenges including those common to cross validation and those common to ilr analysis including the nested dependence of variables. Answering these questions can allow widespread use of phylofactorization for comparison across datasets, annotations on the tree of life, and development of disease detection or, more broadly, community classification despite novel or non-overlapping species.

Phylofactorization can be implemented in the R package `phylofactor`. To encourage future progress on these challenges, some key functions have been constructed to assist future research along these lines. The R package is available on GitHub at https://github.com/reptalex/phylofactor, and novel methods for choosing numbers of factors, calculating quantiles for null distributions of test statistics, and cross-validation can be added to the R package to improve the robustness of the tool.

### 5.1 Criterion for choosing the number of factors

How many factors should we choose? How many traits can we confidently say are contributing to observed patterns of community composition? We can perform phylofactorization for *D* -1 iterations to construct a full basis, but, for big datasets such as the Earth Microbiome Project [Gilbert et al., 2010] containing hundreds of thousands of species and hundreds of thousands of samples, phylofactorization can be prohibitively expensive, even with appropriate parallelization. Even where computation time is not a limiting reagent, downstream analysis and interpretation of phylofactorization can be time intensive. Choosing the number of factors requires constructing an algorithm that stops phylofactorization once there appears to be no more signal in the data, thereby reducing computational time and saving researchers from having to hypothesize putative traits along insignificant edges in the phylogeny likely to be false positives.

The original paper proposed a stopping function for regression phylofactorization based on a Kolmogorov-Smirnov test of the uniformity of the distribution of *P*- values resulting from *F* tests, referred to as the KS stopping criterion. The justification for the KS stopping criterion was that phylofactorization, at each iteration, is often obtaining test statistics for each edge and, if the null hypothesis is true, the *P*-values from multiple hypothesis tests should be uniform. The KS stopping criterion was demonstrated to allow a conservative stopping criterion for simulations of up to 10 clades with randomly assigned geometric changes in one of two environments. The original test did not specify an alternative hypothesis for the KS test, but this author has found the alternative hypothesis that empirical distribution of P values is greater than the uniform null distribution is more robust; two sided KS tests can lead to missed stops as the P value distribution shifts from right skewed to left skewed in a single factor, and downstream factors cause the P value distribu tion to become even more left skewed; missed stops can lead to a full phylofactorization, which can be prohibitively costly for large datasets.

The KS stopping criterion performs well, but other algorithms could outperform it and simultaneously obtain more accurate statements of certainty about the number of factors. There are many criteria for choosing the number of factors in factor analysis which may motivate robust stopping criteria for phylofactorization. Here, we’ll build on the previous literature by comparing the KS stopping criterion to a popular alternative in factor analysis. Horn’s parallel analysis [Horn, 1965] chooses the number of factors based on simulation of the mean, null distribution of eigenvalues of the correlation matrix for standard Gaussian data. The eigenvalues are ranked from largest to smallest and the researcher retains factors whose variances are greater than the similar ranked eigenvalues from the null correlation matrix.

As an analog of Horn parallel analysis, we perform regression phylofactorization on null datasets comprised of i.i.d. standard log normally distributed abundances. Specifically, 300 datasets *X* were simulated with *D* = 32, *n* = 10 and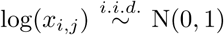. For each dataset, one corresponding independent variable, ***z***, was simulated where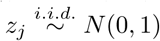. Log normally distributed abundances were used, as opposed to compositional abundances drawn from a logistic normal, for ease (the resulting balances are Gaussian either way) and because for each sample *j*, with total abundance *C*_*j*_= Σ_*i*_*x*_*i,j*_, the isometric log ratios are invariant to *C*_*j*_,

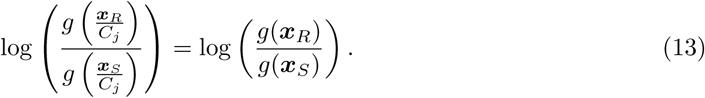

A single, random phylogeny with *D* = 32 species was simulated using the function `rtree` from the R package `ape` [Paradis et al., 2004] and used across all 300 replicate simulations. To simulate non null datasets, we added an association between ***z*** and a set of *b* clades for *b ∈* { 2, 4, 8, 16}.Clades were drawn at random by considering nodes in the tree including the tips, not including the root and its first daughter to ensure *b* correspond to unique factors (the the entire community descends from the root, and the sets of descendants of each of the root’s two daughters are each other’s complement set compositional increases in one are equivalently modeled as compositional decreases in the other). Effects, *β*, were drawn at random from { 3.1, 4.1,…, 18.1 } without replacement the 0.1 included to ensure effects were not an integer multiple of one another as such effects can cancel each other out exactly and are improbable in nature. The sign of association was drawn at random from *α* = {–1, 1}, and the effect was simulated by first simulating null datasets, *X*, as described above, and then, for each clade *c*, with corresponding effects *β*_*c*_, the abundances of all descendants were reassigned as

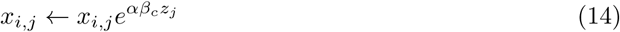

for all *j ∈ c*.

The results of our simulations and comparison are visualized in the first row of Figure 3. The explained variance (EV) for each factor, averaged across all 300 replicates, exhibits a sharp decrease near the true number of factors, *b*. The EV from log normal null datasets exhibits a steady decay until the last factors, where the rate of decay increases. The average null EV curve appears to cross the empirical EV curve at or slightly after the true number of factors. We use this result to construct a “ LN” stopping criterion: a conservative number of factors to include is one less than the first factor at which the average null EV curve is greater than the empirical EV curve.

**Figure 3:**
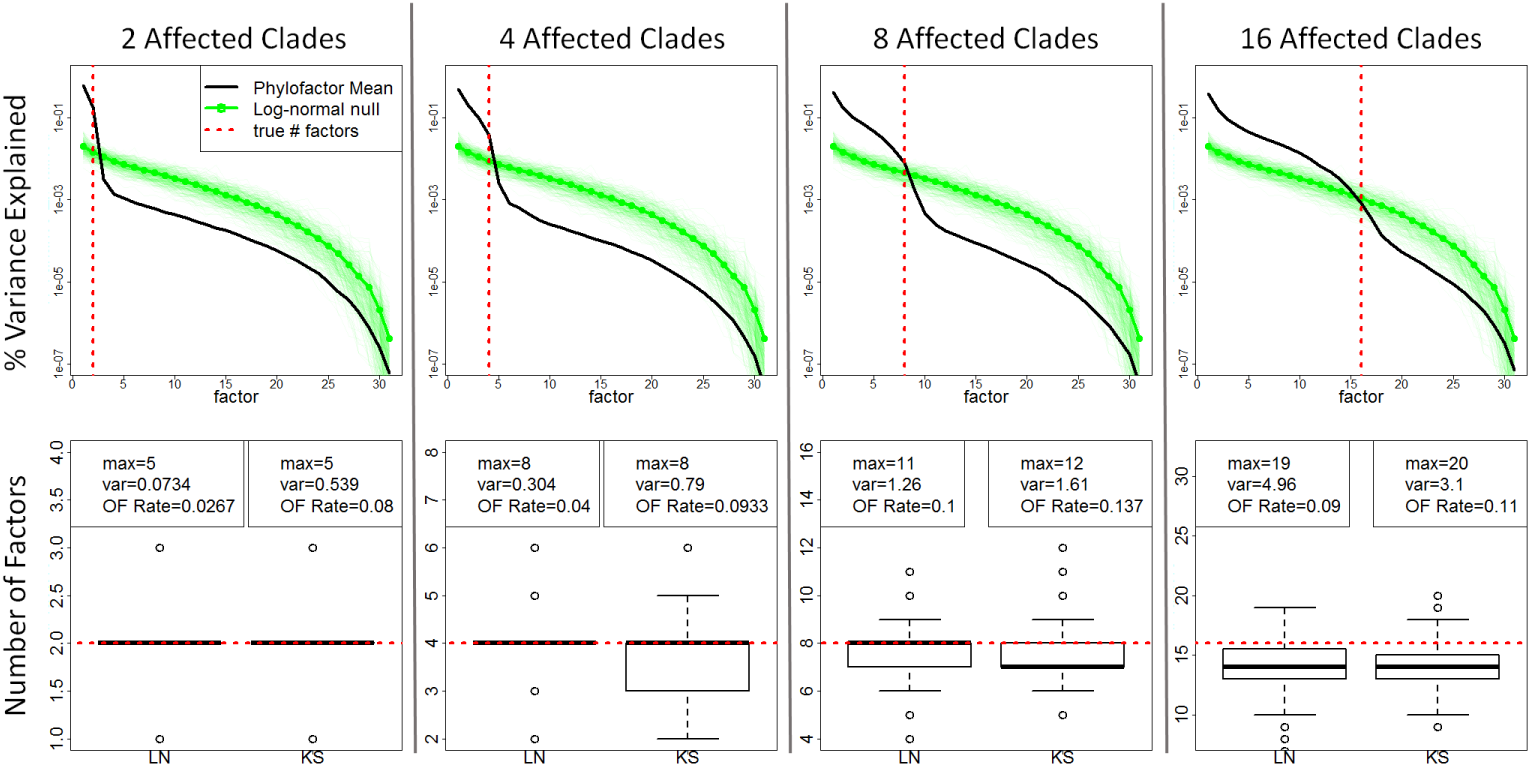
Performance of LN and KS stopping criteria. A challenge of phylofactorization is determining the number of factors, *K*, to include in an analysis, and stopping criteria aim to stop the computationally intensive iteration of phylofactorization. Abundances of *D* = 32 species across *n* = 10 samples were simulated as i.i.d. standard log normal random variables. To simulate effects, a set of *b* clades were associated with environmental meta-data, ***z***, where 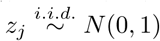. Regression-phylofactorization was performed on 300 dataset for each *b* ∈ {2, 4, 8, 16} and for data with and without effects, with objective function *ω*_ExVar_. (**top row**) The percent explained variance (EV) decreases with factor, *k*, and the mean EV curve for data with *b* affected clades intersects the mean EV curve for null data near where *k* = *b*, motivating a stopping criterion (LN) based on phylofactorization of null datasets to be evaluated and compared to the standard KS stopping criterion. (**bottom row**) The LN stopping criterion has a lower over-factorization (OF) rate than the standard KS stopping criterion (where OF rate is the fraction of the 300 factorizations of data with simulated effects in which *K > b*). Both criteria can be modified to be made more conservative (e.g. the P-value threshold for the KS stopping criterion can be lowered, or the LN criterion can be modified to consider when a quantile of the null EV curves, higher than the mean null EV curve, crosses the empirical EV curve). The KS stopping criterion is far less computationally intensive for large datasets, but the LN criterion may facilitate further statistical statements regarding the null distribution of test statistics in phylofactorization.

To compare the KS stopping criterion with the LN stopping criterion, 300 datasets were simulated with *b* affected clades having an association with meta data, and effects drawn and implemented as described above. Two phylofactorizations were implemented, one in which the phylofactorization continued until the *P* value from the KS test exceeded *P* = 0.01, and another in which phylofactorization continued until the empirical EV was less than the average null EV. In both cases, the factor which satisfied the stopping criterion (*P* > 0.01 for KS, *EV*_*null*_> *EV*_*obs*_ for LN) was not included. The empirically observed number of factors, *b*^*^, for each stopping criterion was compared to the true number of affected clades, *b*.

The performances of the different stopping criteria are visualized in the second row of Figure 3. The KS stopping criterion has a higher over-fitting rate than the LN, especially for low numbers of affected clades. Such high over-fitting rate could be remedied by a lower *P*- value threshold for the KS test. However, the KS stopping criterion is much less computationally intensive-it doesn’t require null phylofactorizations which can be prohibitively costly for large datasets. The performance gains from our analog of Horn’s parallel analysis may not outweigh the computational costs, especially for large datasets (a microbiome dataset of intermediate size may have 1, 000 species and dozens of samples). An analytical solution for the mean EV curves under phylofactorization of standard log-normal data may prove less computationally intensive and yield a more robust stopping criterion than the KS stopping criterion. Other stopping criteria may also consider using rank estimation of *X* to bound criteria for choosing the number of factors. To assist future research in this area, the R package `phylofactor` contains a function `pf.nullsim` to simulate null phylo factorization of log normally distributed data, as done here, for comparison with future stopping criteria.

A related problem to the criteria for choosing the number of factors is the more general problem of null distributions of test-statistics under phylofactorization. For a given factor, what is the probability of an objective function as large or larger than the one observed? For a Bayesian analysis, how does an observed objective function, such as a large explained variance in regression phylofactorization, change a prior distribution of effects on edges in the phylogeny? Phylofactor ization aims to be an inferential tool making inferences on edges in the phylogeny, but, due to the complexity of statistical inference in phylofactorization, currently it is mostly an exploratory tool. Analytical derivations or estimations of the null distribution of test statistics from phylofactoriza tion, in particular for those from regression phylofactorization, can move phylofactorization from an exploratory tool to a more grounded, inferential tool.

For example, consider the F-statistic from regression on 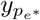for a factored edge, *e*^*\**^, from regression phylofactorization. Under a generalized linear model of null data like Equation (4), the F statistic will follow an F distribution with degrees of freedom *d*_1_= *m* and *d*_2_= *n*- 1*-m*. However, regression phylofactorization on a tree with no polytomies and an objective function to maximize the F statistic will choose, *F*_*max*_, the largest of 2*D*- 3 F statistics. If the F statistics were independent and identically distributed, we could use the fact that

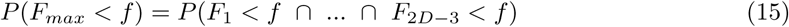

to calculate *P* (*F*_*max*_*< f*) = *P* (*F*_*i*_*< f*)^2*D*-3^. If the F statistics were independent, the problem of the null distribution of regression phylofactorization F statistics would be solved.

Such approximation works well for the early factors of large trees but the approximation worsens for small trees and later factors of large trees due to an increasing percent-overlap of the groups, { *R*_*e*_, *S*_*e*_}, for each edge causing increased dependence among the F statistics, *F*_*i*_. There is a need for more accurate and universal approximations that are robust to tree size and factor number. Such approximations are crucial for disease detection and community classification, for which accurate false positive rates are necessary for clinical use.

As with stopping criteria, phylofactorization of null datasets can be used to produce approximate statements of significance where such statements are desired. Alternatively, if one is interested in the significance of a particular factor, *k*, one can obtain the sub-graphs considered at that level of factorization, the corresponding groups *R*_*e*_ and *S*_*e*_, and simulation of the null distribution-by constructing null ilr balances using log-normal data as above-may yield accurate null distributions. In anticipation of such uses, the R package `phylofactor` contains a function `pf.getGroups` to obtain the set of groups, {*R*_*e*_, *S*_*e*_}, considered at a given iteration of phylofactorization. Analytical tools, or computational tools that don’t require repeated phylofactorization (e.g. direct, null simulation of test-statistics under the dependence observed in the phylogeny), approximating the null distribution of test-statistics and the likelihood functions of parameters (e.g. *β* from regression) given observed test statistics can greatly improve phylofactorization as an inferential tool and allow Bayesian phylofactorization with priors over edges and their associated effects.

Future work on stopping criteria and null distributions of test statistics may allow researchers to make more accurate statements about their uncertainty in the quantity of factors and their particular locations in the phylogeny. Accurate stopping criteria and null distributions can allow biologists to make more nuanced predictions from phylofactorization. For instance, quantifiable certainty about the number of factors and their locations can allow researchers to compare the complexity of trait habitat associations tested; if a treatment affects only two or three clades out of 2, 000, further research understanding the affect of treatments on microbes need only focus downstream physiological studies on two or three clades. Management of a community through such a treatment, as one may hope to modulate communities for improved human health, may be allow precise interventions to affect a few clades. However, if a treatment affects hundreds of clades, and that number *S*^*\**^ can be claimed to be significantly higher than another, management of a community through treatment becomes more complex. Knowing whether a treatment targets 1, 10 or 100 clades, and knowing how uncertain we are about the effects on each clade, can allow rapid progress on modulation of microbial communities.

### 5.2 Cross-validation

If we find mammals are more abundant than reptiles in high, northern latitudes (the mammal/reptile ratio increases as we move North), we can easily probe the generality of this result by cross-validating our finding in southern latitudes. We can easily cross-validate studies in macroscopic ecology because we can readily identify members of the same clades across environments (there is a clear definition of what is a ‘mammal’ and what is a ‘reptile’). Cross validation of findings in microbial communities, for which most of the phylogeny is not annotated, requires consideration of the logic underlying phylogenetic cross validation. Phylofactorization makes inferences that correspond to edges, and consequently can permit cross validation across studies, including those with disjoint sets of species.

The idea behind cross validation of phylofactorization is that a set of *K* edges or chains of edges 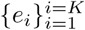 are identified in a dataset ***X***_1_ with phylogeny *T*_1_, and a subset *A ⊆* {1*, …, K*} of those edges,{*e*_*i,*1_}_*i∈A*_, are assessed for their agreement in a second dataset, ***X***_2_, with a phylogeny *T*_2_. The species composition of *T*_1_ and *T*_2_ can range from identical to disjoint, and both *T*_1_ and *T*_2_ are sub-graphs of a universal phylogeny, *T*_*U*_. Cross-validation could be between two datasets, or it could be implemented repeatedly between two subsets of one dataset to prevent over fitting. The challenges of cross validation of phylofactorization are both topological and statistical.

The topological challenges of cross validation require translating the logic behind cross validation of phylogenetic inferences, such as the mammal/reptile comparison above, onto graph topology. How can we compare mammals/reptiles in Australia with the same in North America, when the sets of species are completely disjoint? By reliable identification of the clades, ‘mammals’ and ‘reptiles’, i.e. the groups, *R*_*e*_ and *S*_*e*_ in phylofactorization. Two major issues arise: interruptions of factored edges and whether to include previously identified factors, which implies downstream edges contrast sub graphs, or not, which may make cross validation of downstream edges more robust to erroneous or irrelevant edges in previous factors. These challenges are illustrated in Figure 4.

**Figure 4:**
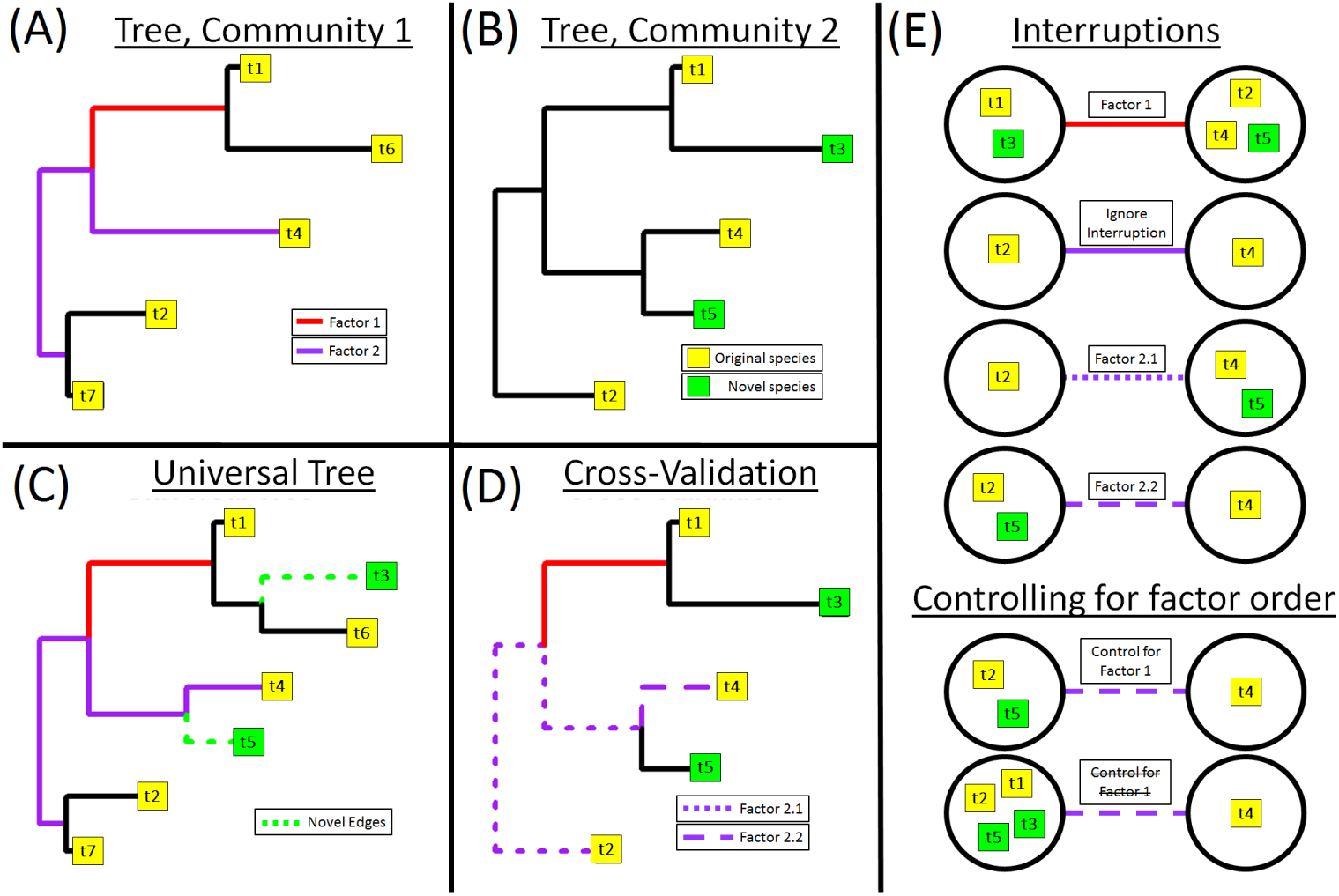
Topological considerations for cross validation. Phylofactorization constructs coordinates that can be used to compare datasets and cross-validate findings, but comparisons require consideration of the graph-topology underlying phylofactorization. **(A)** One community with two factors. The second factor forms a partition separating t4 from {t2,t7}. The second factor does not correspond to a single edge, but instead a chain of two edges. **(B)** A second community, missing species t6 and t7, but containing novel species t3 and t5. **(C)** All factors can be mapped to chains of edges on a universal phylogeny. Novel species “interrupt” edges in the original tree, and cross validation requires deciding what to do with novel species and interrupted edges. Species t3 does not interrupt a factored edge, and so t3 can be reliably grouped with t1 in factor 1. However, species t5 interrupts one of the edges in the edge path of factor 2. **(D E)** Interruptions can be ignored, or they can be used to refine the location of important edges (illustrated in Factor 2.1 and Factor 2.2). Another topological and statistical question is when/whether it is most appropriate to control for factor order. For instance, controlling for factor order with Factor 2.2 would partition t4 from {t2,t5}. Not controlling for factor order would partition t4 from {t1,t2,t3,t5}. Controlling for factor order allows direct correspondence between phylofactorization across datasets, but it may cause later factors to be sensitive to errors with earlier factors, which may be false positives in the original dataset or irrelevant in novel datasets. Two applications for robust cross validation of phylofactorization are disease diagnosis (e.g. Crohn’s disease) and annotation of the universal phylogeny to allow prediction environmental associations of novel microbes.

Interruptions are nodes present in *T*_*U*_ and *T*_2_ which were not present in *T*_1_. For instance, comparing birds to non bird reptiles in Europe (with phylogeny *T*_1_) is contrasting taxa along a single edge in *T*_1_; such a comparison in Australia (with phylogeny *T*_2_) would be complicated by the presence of crocodiles, whose lineage “interrupts” the edge in between the most recent common ancestor of birds and the most recent common ancestor of birds and lizards in Europe. There are two options for how to deal with interruptions: one could either ignore the interruption (remove all descendants of the interrupting node from downstream analysis), or use the interruption to refine the location of the phylogenetic factor (does the defining feature of birds come before or after crocodiles?). Ignoring interruptions allows direct comparison of groups which led to the inference of a factor in the original dataset, but refining the location of a phylogenetic factor may increase the power across datasets by increasing the number of taxa and allowing more focused downstream studies of microbial physiology. The R package `phylofactor` contains a function `crossVmap`, which takes as input the groups, {*R*_*e*1_, *S*_*e*1_} and the universal tree *T*_*U*_, and inputs either the groups {*R*_*e*2_, *S*_*e*2_} corresponding to the direct comparison (ignoring interruptions), or the set of all possible groups {*R*_*e*2,*i*_, *S*_*e*2,*i*_} corresponding to each edge in the link of edges to permit refinement of the location of the phylogenetic factor (e.g. one can obtain the set of groups, { birds + crocodiles, reptiles } and { birds, lizards+ crocodiles}, to determine where the meaningful difference between birds and reptiles arose).

The second topological challenge is how to deal with the hierarchical structure of factors. Given an ordered set of factored edges 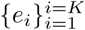, should one cross validate the edges in the order in which they were factored for the original dataset, or should one perform some analog of phylofactorization on this set of edges? The former allows a strict comparison of phylogenetic factorization across datasets, whereas the latter may be less sensitive to errors in the exact sequence of factors and the presence of erroneous edges or edges corresponding to traits with functions in one community but not in others.

The statistical challenges of cross-validation include standard challenges of compositional data analysis (e.g. the nested dependence of pre-determined ilr balances) and particular challenges to statistical analysis under different topological considerations listed above. Many of these challenges may have easy solutions already in existence but unknown to this author. Robust cross validation within datasets, by repeated training and testing of subsets of the same dataset, can reduce over fitting (and thereby be related to the criteria for determining the number of factors), and robust, one time cross validation between two datasets can allow phylogenetically informed diagnostics of microbial community state. For example, if one finds phylogenetic factors driving Crohn’s disease [Kostic et al., 2014], it may be possible to develop diagnostic tools to assign the likelihood a patient has Crohn’s disease given a sample of their intestinal microbial communities, even if their intestinal microbial communities have novel species and interrupting nodes. For another example, cross validation can allow refinement of the location of edges and allow for annotation of a universal tree of life to allow for prediction of environmental associations of novel, uncultivated microbes.

## 6 Discussion

Communities are compositions of species, and species have traits, evolved by natural selection, which determine their relative abundances, habitat associations and responses to perturbation. Traits can be mapped to edges on the phylogeny, and consequently one can describe communities as the species or as mixtures of traits, latent variables defined through phylogenetic coordinates. Phylofactorization is a method for choosing phylogenetic coordinates along which communities change, coordinates whose positions are interpretable as latent variables or functional ecological traits that evolved along the phylogeny. The phylogeny is a graph, and phylofactorization is a graph-partitioning algorithm with no balance constraint which iteratively chooses edges of importance and then partitions the phylogeny along that edge. Often, to choose which edge to cut, one is interested in a measure of difference between the two groups, as partitioning the graph along edges that “best differentiate” two groups may result in a set of sub-graphs with low within group differences and “differences” can be the presence/absence of a trait. The balancing elements from the isometric log ratio transform serve as a standardized measure of difference and a means to construct an orthonormal basis from phylofactorization.

An important special case of phylofactorization is regression phylofactorization. Because communities can often be justifiably analyzed as compositions, especially microbial communities sampled through amplicon sequencing, regression phylofactorization entertains balances corresponding to a partition along each edge and partitions the graph along whichever edge maximizes an objective function for regression. Regression phylofactorization can be interpreted as hierarchical regression, construction of a decision tree about species’ associations with meta data, and a latent variable model whose latent variables are traits, with similar challenges to factor analysis (although, strictly speaking, it is not necessarily factor analysis). By constructing a sequential binary partition corresponding to a sequence of edges in the phylogeny which explain the most total variance in a compositional dataset, regression phylofactorization constructs a low rank approximation of a dataset with orthonormal “loadings” (balancing elements) whose balances correspond to the relative abundances of organisms with and without a putative trait, controlling for other, previously identified putative traits.

Much like PCA, factor analysis and other clustering algorithms in their initial conception, phylo factorization is a predominantly exploratory tool, but future work on criteria for choosing the number of factors and the null distributions of test statistics can assist the use of phylofactorization for rigorous statistical inference on edges in the phylogeny and associations between clade abundances and meta data. Significant progress towards understanding microbes and their traits can be made with robust cross validation of phylofactorization to allow comparison of inferences across datasets, including datasets containing novel species. Cross validation of phylofactorization requires careful consideration of the graph topological inferences being made, but such tools can permit novel methods for disease detection and refined inferences on a universal phylogeny to allow predictions of habitat associations of novel, uncultivated microbes.

The challenges listed above are a small subset of future research directions which may collectively bring about more accurate, inferential phylofactorization and greatly accelerate research in the microbiome world for which most of the phylogeny is unannotated and most species have never been observed under a microscope. There are many other challenges not listed here but which are still important. For instance, like all methods in compositional data analysis, there is a need to understand the appropriate treatment of zeros for phylofactorization, which may depend on the underlying model of count data. For some ecological effects there may be more appropriate contrasts of two groups split by a partition, such as arcsines of differences of total relative abundance in two groups, *R* and *S*, used for testing of neutral drift in ecological time series [Washburne et al., 2016]. To assist future research on phylofactorization, the R package `phylofactor` has several functions aimed at providing user interface with internal objects for method development. The R package is available at https://github.com/reptalex/phylofactor.

## Acknowledgments

Phylofactorization was conceived under the support of Diana Nemergut, who passed away as the method was being developed by this author. Under support from D. Nemergut’s start up funds, generously made available by Duke University, the method was further nourished by the networking, support and fruitful discussions with Juanjo Egozcue, Vera Pawlowsky Glahn, Rob Knight, Jamie Morton, Noah Fierer, Justin Silverman and Lawrence David.

## Appendix: Mathematical Notation

- *D*: number of species in a community.
- *n*: number of communities sampled in a dataset.
- *m*: number of meta data variables
- *p*: partition index. Each partition defines two disjoint index sets over the species in the community, {*R*_*p*_, *S*_*p*_}. *p*_*e*_ denotes partition defined by a given edge, *e*
- *p|k*: generic partition of one remaining group after *k* previous partitions.
- *e*: edge index. In this paper, edges will often have corresponding partitions dependent on the sub graphs in which the edge is found.
- Δ^*D*-1^: Simplex {***x*** *∈* ℝ^*D*^*| x*_*i*_*≥* 0 *∀i* and ∑*i x*_*i*_= 1}.
- ***x***_*j*_: compositional vector in sample *j*, ***x*** *∈* Δ^*D*-1^.
- *C*_*j*_: total abundance of a non-normalized abundance vector, ∑_*i*_ *x*_*i, j*_.
- *𝒱*_*p*_: balancing element corresponding to partition, *p*.
- *𝒱*_*p|k*_: balancing element orthogonal to *k* previously defined balancing elements 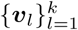
- *y*_*p,j*_: ilr transform, a.k.a. balance, of sample *j* determined by partition, *p*. Used to denote candidate balances when the full sequential binary partition is not yet chosen.
- ***z***_*j*_: *m* vector of environmental meta-data in sample *j*.
- ***X***: Data matrix, (*x*_*i,j*_) *∈* Δ^*D*-1*×n*^, whose rows are species and columns are compositional vectors, ***x***_*j*_.
- *R*_*p*_: index set of species in one of two, disjoint groups defined in a partition. Absence of subscript implies a generic partition, sometimes subscript *e* is used as shorthand to denote partition defined by a given edge, *p*_*e*_.
- *S*_*p*_: index set of species in the complementary group to *R*_*p*_, i.e. the other group defined in a particular partition.
- *g*(***x***): geometric mean of ***x***.
- clr_*i*_(***x***): *i*th element of centered log ratio transform, clr_*i*_(***x***) = *xi/g*(***x***).
- ilr_*i*_(***x***): *i*th element of a given isometric log ratio transform (related to *y*_*i*_, but corresponding to a complete transform instead of a candidate ilr balance).
- clr(*X*), ilr(*X*): clr or ilr-transformed dataset. In this paper, these operations are applied to columns of *X*.
- *ω* arbitrary objective function.
- *ω*_𝕧_:variance of ilr balance
- *ω*_ExVar_: explained variance from regression on ilr balance

